# Sex-biased gene expression shapes sex differences in gene essentiality

**DOI:** 10.64898/2026.04.13.717533

**Authors:** Clarissa Rocca, Alex R. DeCasien

## Abstract

Sex differences in disease incidence and progression are well documented, yet their underlying molecular mechanisms remain poorly understood. Multiple models suggest that baseline gene expression levels shape the impact of gene disruption, raising the possibility that sex-biased expression itself contributes to sex differences in cellular vulnerability. Here, we test this hypothesis by integrating sex-biased transcriptomic profiles with large-scale CRISPR loss-of-function screens to determine whether sex-biased expression predicts sex-biased gene essentiality across the genome. We find that gene expression level and sex chromosome dosage each explain a modest fraction of variance in essentiality, with substantially larger effects for sex chromosome genes than for autosomes. Across genes, sex effects on expression and essentiality are small in magnitude but directionally aligned, suggesting that sex differences in transcription can influence functional dependency. To resolve how these relationships arise, we applied gene-level mediation analyses to decompose sex effects on essentiality into expression-mediated and expression-independent components. This approach revealed multiple mechanistic architectures. On autosomes, most genes exhibited either sex-biased essentiality from direct sex effects (independent of expression) or sex-biased expression without functional consequence, while expression-mediated sex differences accounted for a smaller but substantial fraction of genes. In contrast, X chromosome genes were dominated by direct, expression-independent sex differences, consistent with strong effects of sex chromosome dosage, but also showed enrichment of expression-mediated architectures, particularly among X gametologs. Together, our results demonstrate that while sex-biased expression can generate sex-biased gene essentiality, this mechanism is not the default. Instead, sex-biased functional dependency is often driven by direct, expression-independent effects, particularly on the X chromosome, where dosage and compensatory mechanisms play a dominant role.

## Introduction

Sex differences in the incidence, presentation, and progression of many diseases are well documented [1–3], yet the molecular mechanisms underlying these differences remain incompletely understood (Note 1). One proposed explanation is that males and females differ in their baseline gene expression profiles, creating distinct molecular environments that shape how genetic perturbations affect cellular function.

A growing body of work supports the idea that baseline expression levels modulate the consequences of genetic variation and gene disruption. However, existing models make differing predictions about how expression influences functional outcomes. Regulatory-threshold models suggest that sex differences in disease risk arise not from sex-biased expression of the causal genes themselves, but from broader sex-biased biological pathways that shape how genetic variants act within cells and tissues. In this framework, males may occupy, for example, a baseline regulatory state that lies closer to autism spectrum disorder (ASD) associated expression thresholds, making them more susceptible to regulatory variants that shift expression of these pathways into a pathogenic range [4–6]. Studies of coding variation suggest contrasting roles for expression: work on partial loss-of-function variants in ASD indicates that higher expression may buffer females against damaging coding variants, reducing their phenotypic effects when gene dosage is partially compromised [7], whereas studies of haplotype-dependent penetrance suggest that higher expression can instead amplify the impact of damaging variants by increasing the dosage of mutant transcript [8]. More generally, essential genes tend to be more highly expressed than nonessential genes, underscoring a broader link between expression level and the functional consequences of gene disruption [9–12].

Together, these observations suggest a general principle: the impact of gene disruption depends on the expression context in which it occurs. If males and females differ in baseline expression levels, then the same perturbation could produce systematically different functional outcomes. This raises the possibility that sex-biased gene expression contributes to sex-biased gene essentiality – that is, differences in the requirement of genes for cell survival between sexes.

Here, we test this hypothesis by integrating sex-biased expression profiles [13] with large-scale CRISPR loss-of-function (LoF) screens [14] from the Achilles project to assess whether sex-biased expression predicts corresponding sex-biases in gene essentiality. In this context, gene essentiality measures the functional requirement of a gene for cell survival; a high essentiality score indicates that its loss-of-function compromises cell viability. Although prior work using these data demonstrated that sex chromosome dosage affects both gene expression and essentiality [13], it did not integrate these two measures to test for a causal link. Furthermore, the relationship between sex-biased expression and essentiality may vary across genomic contexts (e.g., autosomal vs. X chromosome genes) and functional categories (e.g., broadly expressed vs. testis-elevated genes).

Motivated by these gaps, we asked three questions. First, how much of the variance in gene essentiality can be attributed to gene expression level and sex chromosome dosage? Second, does sex chromosome dosage have concordant effects on gene expression and essentiality across genes? Third, does gene expression mediate the relationship between sex chromosome dosage and essentiality? To address these questions, we analyze gene-level effects across multiple dimensions of sex chromosome variation, including sex (XX vs. XY), X chromosome dosage (XX vs. X0), and Y chromosome dosage (Y+ vs. Y−) (Figure 1A). We combine correlation-based analyses with gene-level mediation models to distinguish expression-mediated effects from expression-independent effects on gene essentiality (Figure 1A).

**Figure 1.**
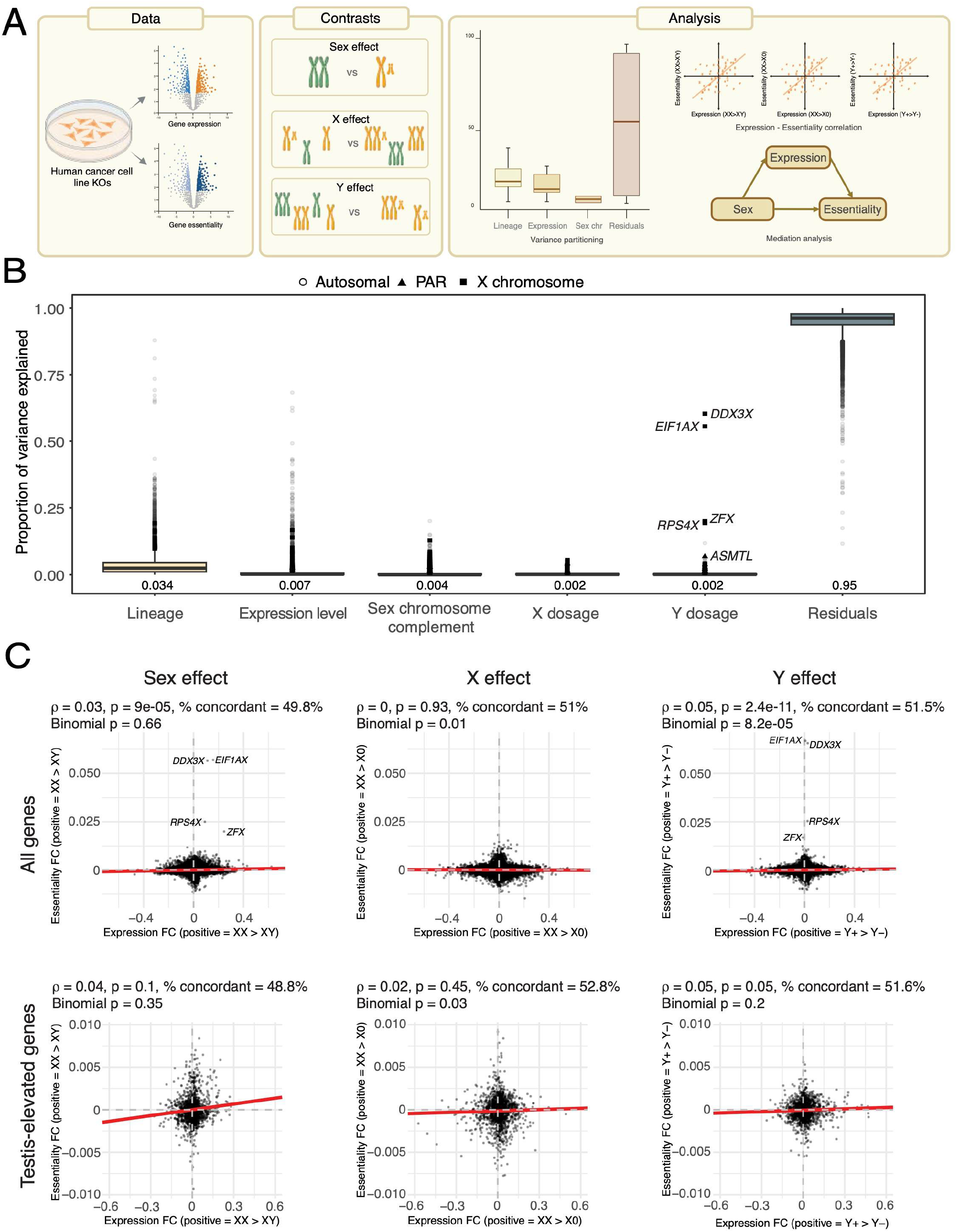
Analytical framework and genome-wide relationships between sex-biased expression and gene essentiality. (A) Workflow used in this study. Gene-level expression and essentiality data were obtained from human cancer cell lines. Analyses were performed across three contrasts: sex (XX vs XY), X chromosome dosage (XX vs X0), and Y chromosome dosage (Y+ vs Y-). Chromosomal complements from females are represented in orange, and in green for males. We used different approaches to analyze this data. Left: variance partitioning to quantify the proportion of variance in essentiality explained by different contributors. Top right: correlation analyses across expression and essentiality datasets. Bottom right: Mediation analysis decomposes sex effect on gene essentiality onto expression-mediated (indirect) and expression independent (direct) components. (B) Boxplots show the proportion of variance in gene essentiality explained by each component across genes. Residual variance represents unexplained variation. Each point corresponds to an individual gene. Circles represent autosomal genes, triangles PAR genes, and squares represent X chromosome genes. (C) Scatterplots showing the relationships between the effects of sex, X chromosome, or Y chromosome on gene expression (x-axes) versus essentiality (y-axes). Each point represents a single gene. Effects are represented on both axes by fold changes (FC). Positive FC values represent higher gene expression or essentiality levels in XX vs. XY (left column), XX vs X0 (middle column), and Y+ vs. Y- (right column). Negative FC values represent higher gene expression or essentiality in XY vs XX (left column), X0 vs XX (middle column), and Y-vs Y+ (right column). Analyses were conducted across all expressed genes (N=17,904) (top row) and testis elevated genes (bottom row). Red lines indicate the best-fit linear regression. Grey dashed lines mark zero (i.e., no effect) on both axes. Subtitles in each plot report Spearman’s ρ, the corresponding p-value, the percent concordance (i.e., % of points in the top right + bottom left quadrants), and the binomial p value of genes in the top right and bottom left quadrants of each plot.

This framework allowed us to directly test whether sex-biased expression provides a mechanistic basis for sex-biased functional dependency across the genome, and to determine the relative contribution of expression-mediated and expression-independent mechanisms.

## Methods

### Data Sources

For our correlation-based analyses (see ‘Statistical analyses’ section below), we obtained results from Shohat and colleagues [13]. Specifically, Supplementary Table 2 in [13] provided gene-level expression log□ fold changes (FCs) and false discovery rates (FDRs), and Supplementary Table 3 in [13] provided gene-level essentiality fold changes (log□) and FDRs. Both datasets included gene identifiers (HGNC symbols), chromosome annotation, pseudoautosomal region (PAR) status, and X-chromosome inactivation (XCI) status. A brief summary of their methods are provided here. In the original study, Shohat and colleagues [13] analyzed 839 human cancer cell lines (371 female-derived and 468 male-derived) using data from the Broad Institute’s Achilles project [14], the largest systematic survey of human gene essentiality. Gene essentiality scores from these screens were modeled using linear mixed-effects models. The models included sex and the number of X and Y chromosomes as fixed effects, with tissue of origin treated as a random effect, thereby enabling separation of the sex effect (estimated from the XX vs. XY contrast), X-chromosome dosage effect (XX vs. X0 contrast), and Y-chromosome dosage effect (Y+ vs. Y- contrast). We directly adopted those contrasts in this study. Throughout, ‘contrast’ refers to the pairwise comparison (e.g., XX vs. XY); ‘effect’ refers to the estimated difference between groups from the linear models; and *‘*fold change (FC)’ refers to the log□-transformed ratio of expression or essentiality values between groups. For mediation analyses, we used gene-level CRISPR-Cas9 essentiality scores and bulk RNA-seq expression profiles from DepMap version 22Q1[15], together with sex chromosome complement and inferred sex chromosomes information from Shohat and colleagues Supplementary Table 1 [13].

### Variance partitioning

To quantify the contribution of gene expression, sex chromosome complement (SCC), X dosage, Y dosage, and tissue of origin to variability in gene essentiality, we performed variance partitioning using the *variancePartition* R package [16]. Cell lines lacking a lineage annotation were excluded (N=1). Age at sampling was not included in the analysis due to the elevated number of missing data. For each gene, we fit a mixed model with gene expression, X chromosome copy, and Y chromosome copy as fixed-effects and sex chromosome complement and lineage as random effects. Specifically, for each gene *g*, we modeled essentiality as: *ess*_*g*_ ∼ expr_*g*_ *+ X_copy + Y_copy + (1*|*SCC) + (1*|*Lineage)*. Models were fitted by restricted maximum likelihood (REML) using the *lmer* function from the lme4 R package [17]. For each fitted model, variance components for the random effects and the residual error were obtained from the *VarCorr* function. To estimate the contribution of each fixed effect, we computed the variance of the fitted fixed-effect term across samples. The total variance for each gene was then defined as the sum of these fixed-effect variances, the random-effect variances, and the residual variance. Variance proportions were calculated by dividing each component by the total. We summarized variance proportions across genes by taking column medians of the per-gene variance fraction matrix, and visualized the distributions of variance fractions using boxplots of the melted per-gene variance fraction output.

### Gene sets

For each contrast, we tested the relationship between expression and essentiality across multiple subsets of genes. These included subsets based on chromosomal location: all genes, autosomal genes, and X chromosome genes (defined in the original publication [13]). We also examined a subset of genes that escape X chromosome inactivation (XCI), as they showed dosage-sensitive expression in XX cell lines [13]. In addition, we conducted targeted analyses on: i) genes involved in synaptic transmission (N=432; GO:0007268) and immune response (N=1,775; GO:0006955); ii) genes with elevated expression in sex-specific tissues (N=1,590 testis-elevated; N=178 ovary-elevated) obtained from the Human Protein Atlas [18]; iii) genes implicated in sex-biased conditions, including rare-variants in autism spectrum disorder (ASD) (N=102) [19] - also subdivided into those involved in neuronal communication (N= 58) or gene expression regulation (N=24) [20,21], Parkinson’s disease associated genes derived from Genomics England PanelApp (N=45) [22] (Parkinson Disease and Complex Parkinsonism, high-confidence genes only), and multiple sclerosis associated genes based on OMIM (N=169) [23] annotations (Table S2).

### Mediation analysis

After excluding cell lines with missing sex chromosome complement annotations, we ran mediation analysis on sex (N=175 XX, N=223 XY) and dosage (X or Y effect) (Figure 1A). Using sex chromosome complement categories from Shohat and colleagues [13], we derived integer-valued X_count and Y_count variables for each cell line (e.g. Female_X0: X_count = 1, Y_count = 0; Female_XX: X_count = 2, Y_count = 0; Male_X0: X_count = 1, Y_count = 0; Male_XY: X_count = 1, Y_count = 1; Male_XXY: X_count = 2, Y_count = 1).

We fitted a total of six linear models per gene: two for each of sex, X, and Y effect (*lm* function in the R package *stats[24]*). We used: i) a mediator model (expression ∼ sex or dosage) and ii) an outcome model (expression ∼ sex or dosage + expression). These functions were passed to the *mediate* function from the R package *mediation[25]*. For each gene, we estimated the average causal mediation effect (ACME – indirect effect of sex on essentiality through expression), the average direct effect (ADE – effect of sex on essentiality not operating through expression), the total effect and the proportion mediated, together with 95% confidence intervals obtained from bootstrap resampling (500 simulations per gene). To account for multiple testing across genes, we applied FDR using *p.adjust(method = “fdr)* in R. Chromosomal annotation of genes (X chromosome vs. autosome vs. Y chromosome) was obtained from Ensembl via the *biomaRt* R package [26]. After estimating mediation statistics for each gene, we classified genes into different categories based on nominal significance thresholds (p < 0.05) and the directionality of effects. Significance was defined for the sex effect on expression, the direct effect on essentiality, the total effect on essentiality, and the mediation effect.

We first assigned genes to six mediation patterns based on the presence and sign of significant average direct effects (ADE) and average causal mediated effects (ACME): pure mediation (ACME significant only), direct-only (ADE significant only), consistent mediation (ACME and ADE significant with the same sign), inconsistent mediation (ACME and ADE significant with opposite signs) and no effect (no significant ACME, ADE, or total effect).

We then used these mediation patterns, together with significance of sex effects on expression and essentiality, to define seven higher-level gene categories: (i) mediation-driven sex difference genes, defined by a significant indirect effect (ACME) contributing to sex-biased essentiality, with or without a direct effect; (ii) direct sex difference genes, defined by significant sex effects on essentiality (total effect and/or direct effect) without evidence of mediation (non-significant ACME); (iii) sex-biased expression only genes, defined by significant sex effects on expression without detectable sex effects on essentiality; (iv) no detected sex signal / baseline genes, defined by no significant sex effects on either expression or essentiality; (v) inconsistent mediation genes, defined by significant indirect (ACME) and direct (ADE) effects of opposite sign; and (vi) opposite-sign total versus direct effects genes, defined by significant total and direct effects with opposite signs.

### Statistical analyses

All analyses were carried out in R version 4.5.1 [24].

For each contrast and gene subset, we evaluated the relationship between the expression FC and essentiality FC using both Pearson’s and Spearman’s correlation coefficients (*cor.test* function in the R package *stats [24]*). We also calculated the percentage of genes showing concordant FC directions (e.g., genes located in the upper-right or lower-left quadrants of the expression FC vs essentiality FC scatterplots). To visualize these relationships, we fitted simple linear regression models of the form *essentiality FC ∼ expression FC* to estimate the slope and intercept of the association. Finally, we performed binomial tests (*binom.test* function in the R package *stats [24]*) to determine whether the observed concordance proportion exceeded the expectation under random chance. Unless otherwise specified, analyses were conducted on both signed FC (directional) and absolute FC (magnitude).

### Visualization

Scatterplots of expression FC vs. essentiality FC were generated using *ggplot2 [27]*, including grey dashed zero axes and red regression lines. Subtitles reported Spearman’s ρ, corresponding p-values and concordance; for signed FC analyses, percent concordance was also shown.

## Results

### Variance partitioning reveals limited contributions of expression and sex chromosome dosage to gene essentiality

We first quantified the drivers of gene essentiality across 763 cancer cell lines using variance partitioning (Figures 1A,B). To do so, we modeled gene essentiality as a function of lineage, gene expression level, sex chromosome complement, X dosage, and Y dosage (Methods). Lineage was the largest contributor to variance in essentiality (median = 3.4%) (Figure 1B, Table S3). Gene expression explained a modest fraction of variance (median = 0.7%) (Figure 1B, Table S3), indicating that expression levels account for only a small portion of variability in essentiality. Variance attributable to X and Y chromosome dosage was minimal for most genes (median = 0.2% for each), and sex chromosome complement explained little variance overall (median = 0.4%) (Figure 1B, Table S3). Most gene-level variability in essentiality was captured by the residual component (median = 95.0%) (Figure 1B, Table S3). Together, these results indicate that while gene expression and sex chromosome dosage contribute to gene essentiality, their effects are modest relative to other sources of variation.

### Sex chromosome dosage effects on gene expression and essentiality are weak but directionally aligned

We next asked whether sex and sex chromosome dosage have concordant effects on gene expression and gene essentiality across genes. To address this, we compared fold changes (FCs) for expression and essentiality across three contrasts: sex (XX vs. XY), X chromosome dosage (XX vs. X0), and Y chromosome dosage (Y+ vs. Y−).

Across all expressed genes present in both datasets (N=17,283 genes), sex differences in gene expression and gene essentiality were weakly but positively correlated (ρ = 0.03, p = 9e-05), with 49.8% of genes showing concordant directionality (Figure 1C; Table S1). Restricting the analysis to genes with detectable effects (FDR < 0.2 in either dataset) increased the correlation (ρ = 0.09, p = 0.01) and concordance (∼58%), indicating that alignment between expression and essentiality is more apparent among genes with stronger effects. These findings indicate that genes more highly expressed in one sex tend to also be more essential in that sex, although the magnitude of this relationship is small.

This relationship was weaker among autosomal genes (ρ = 0.02, p = 0.03, 49.3% concordant) (Table S1), but showed increased directional concordance among X chromosome genes (62.2%), despite no significant correlation in effect size (ρ = 0.01, p = 0.75) (Table S1). Consistent with this, analyses of absolute fold changes for X chromosome genes showed modest but significant associations both across all X-linked genes (ρ = 0.08, p = 0.04) and among X-linked genes passing the FDR threshold (ρ = 0.10, p = 0.03) (Table S1), suggesting limited coupling in effect magnitude even when directional concordance is weak.

We next examined chromosome dosage effects. X chromosome dosage (XX vs. X0) showed little evidence of concordance between effects on gene expression and essentiality (Figure 1C; Table S1). Across autosomal genes, the correlation was minimal (ρ = 0.02, p = 0.02; 51.7% concordant) (Table S1). In contrast, Y chromosome dosage (Y+ vs. Y−) showed the clearest evidence of concordance. Across all genes, Y dosage effects on expression and essentiality were positively correlated (ρ = 0.05, p = 2.38e-11; 51.5% concordant), with similar results for autosomal genes (ρ = 0.06, p = 4.41e-13; 51.5% concordant) (Figure 1C; Table S1). Restricting the analysis to autosomal genes with detectable effects further increased the correlation (ρ = 0.15, p = 1.87e-4) and concordance (55.9%), indicating that genes most responsive to Y chromosome presence show stronger alignment between expression and essentiality.

We further examined these relationships across functionally defined gene sets, genes with elevated expression in sex-specific tissues, genes involved in immune response and synaptic transmission, and genes associated with sex-biased diseases (Tables S1, S2) (Methods). Most gene sets did not show consistent associations between expression and essentiality; however, several subsets showed suggestive patterns (Table S1).

Testis-elevated genes showed a positive correlation for Y chromosome dosage effects across all genes in the set (ρ = 0.05, p = 0.05; 51.6% concordant), consistent with a role for Y-linked regulatory context in shaping male-biased functional programs (Figure 1C; Table S1). Parkinson’s disease-associated genes showed a positive correlation for Y dosage effects across all genes in the set (ρ = 0.32, p = 0.03; 64.4% concordant), with similar results for the autosomal subset (ρ = 0.32, p = 0.03; 62.8% concordant), although these analyses were based on small numbers of genes (N = 45 and N = 43, respectively) (Table S1). Among X-linked genes, ovary-elevated genes showed moderate correlations for sex effects on absolute fold changes (for genes passing FDR threshold; ρ = 0.61, p = 0.01; N = 16) and for X chromosome dosage effects (ρ = 0.51, p = 0.02; N = 20), while immune response genes showed a moderate association for sex effects on absolute fold changes (ρ = 0.41, p = 0.01; N = 37) (Table S1). These results suggest possible context-specific coupling between expression and essentiality in some gene sets, but such patterns were not broadly observed.

Overall, these results suggest that sex chromosome dosage effects on gene expression and essentiality are generally small in magnitude but exhibit weak directional alignment across genes.

### Sex differences in gene essentiality are largely independent of gene expression

To directly test whether sex-biased expression mediates sex differences in gene essentiality, we performed gene-level mediation analyses (Figure 2A, Table S4) (Methods). For each gene, we decomposed the total effect (TE) of sex on essentiality into an expression-mediated component (i.e., the average causal mediated effect; ACME) and an expression-independent component (i.e., the average direct effect; ADE) (Figure 2A).

**Figure 2.**
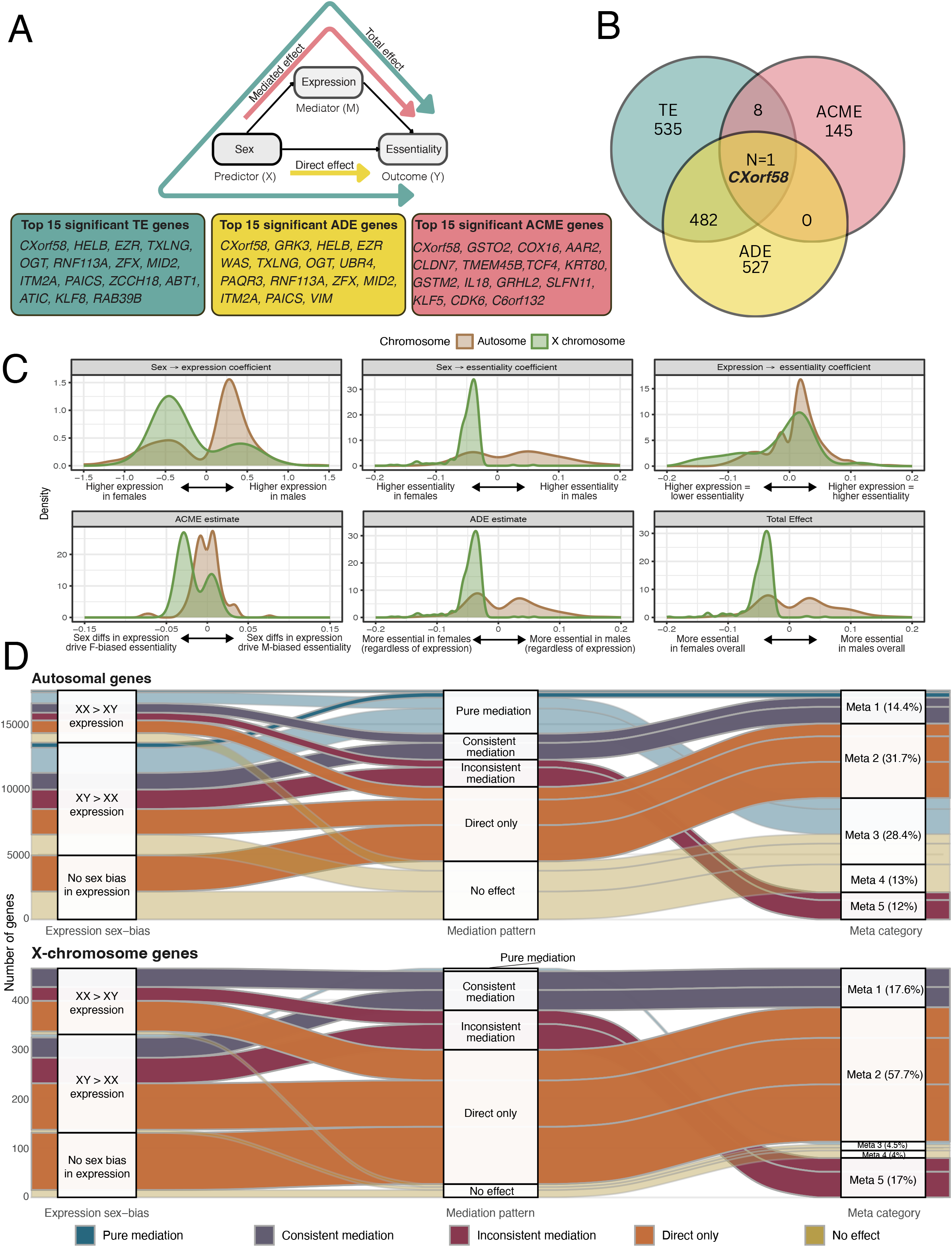
Mediation analysis of sex effects on gene essentiality. (A) Schematic representation of the mediation analysis used to decompose the effect of sex on gene essentiality. Sex is modeled as the predictor (X), gene expression as the mediator (M), and gene essentiality as the outcome (Y). The total effect is indicated by the teal path, the mediated effect by the pink path and the direct effect by the yellow path. Boxes below highlight the top 15 significant genes for each component, ranked by FDR-adjusted p-values: total effect (TE), direct effect (ADE) and mediated effect (ACME). (B) Venn diagram showing the overlap between genes with significant TE in teal,ADE in yellow, and ACME in pink. Only one gene (*CXorf58*) is shared across all three categories. (C) Density distributions of coefficients from mediation analysis shown separately for autosomal genes (brown) and X-chromosome genes (green). Top row: distribution of sex effects on gene expression (left), sex effects on gene essentiality (middle), and the relationship between expression and essentiality (right). Bottom row: distribution mediated effects (ACME), direct effects (ADE), and total effects (TE) of sex on essentiality (mediated by expression). (D) Alluvial plot illustrating the flow of genes from expression sex-bias categories (lect) through mediation patterns (center) to final mechanistic meta-categories (right). Autosomal genes (top) and X-chromosome genes (bottom) are shown separately. Across autosomal genes, the largest fraction of genes falls into the direct-only category (Meta 2, 31.7%), followed by genes with sex-biased expression but no corresponding essentiality effects (Meta 4, 28.4%). X-chromosome genes are strongly enriched for direct-only effect (Meta 2, 57.7%).

Across all expressed genes, most of the significant (FDR-adjusted p < 0.05) sex effects on essentiality were driven by direct, expression-independent effects. Specifically, among genes with a significant total effect of sex on essentiality, the majority also showed a significant direct effect (N = 482), whereas only a small number showed evidence of mediation (N = 8). Only a single gene (*CXorf58*) exhibited significant total, direct, and mediated effects simultaneously (Figure 2B).

Distributions of mediation coefficients revealed that sex effects on expression and essentiality were more pronounced for X chromosome genes than for autosomes, while expression–essentiality relationships were centered near zero (Figure 2C). X chromosome genes were strongly enriched among genes with significant direct and total effects, but not among mediated effects. Genes more essential in females (negative ADE or total effect) were enriched for X-linked genes (ADE: OR = 91.9; p < 2.2e-16; TE: OR = 94.2; p < 2.2e-16) (Table S5).

To further characterize the architecture of sex effects, we classified genes into mechanistic categories based on the presence and direction of direct and mediated effects (Methods). Using nominal significance thresholds (p < 0.05), we classified genes into seven mechanistic architectures based on sex effects on expression, sex effects on essentiality (total and direct), and mediation patterns (ACME). These categories were defined as follows: (i) mediation-driven genes, showing a significant indirect effect (ACME), with or without a direct effect; (ii) direct-only genes, showing significant sex effects on essentiality without evidence of mediation (non-significant ACME); (iii) sex-biased expression only genes, showing significant sex effects on expression without corresponding effects on essentiality; (iv) no detectable sex signal genes, showing no significant sex effects on either expression or essentiality; (v) inconsistent mediation genes, showing significant indirect and direct effects in opposite directions; and (vi) opposite-sign total vs. direct genes, showing significant total and direct effects with opposing signs, consistent with rare suppression or sign-flip architectures.

On autosomes, the most common pattern was a direct sex effect on essentiality independent of expression (31.9%), followed by sex-biased expression without a corresponding effect on essentiality (28.3%) (Figure 2D). Expression-mediated effects accounted for a smaller but substantial fraction of genes (14.1%). Comparable proportions of genes showed no detectable sex bias in either expression or essentiality (13.2%) or exhibited inconsistent mediation (11.9%), in which direct and mediated effects act in opposite directions. Smaller fractions showed unclear mediation despite a sex difference (0.6%) or opposite-sign total vs. direct effects (<0.1%), indicating rare but complex architectures.

In contrast, X chromosome genes showed a markedly different architecture (Figure 2D). A majority exhibited direct, expression-independent sex differences in essentiality (57.0%), consistent with strong effects of sex chromosome dosage and X-specific regulatory context. Some X gametologs were assigned to this category (*NLGN4X, USP9X, EIF1AX*). Expression-mediated sex differences were more common on the X chromosome than on autosomes (19.1% vs. 14.1%), suggesting that dosage-sensitive expression of X-linked genes more frequently contributes to sex-biased dependency. Most X gametologs were assigned to this category (*DDX3X, RPS4X, ZFX, TXLNG, KDM5C, KDM6A*). In addition, inconsistent mediation was more common (17.0% vs. 11.9%), suggesting widespread compensation between expression and direct effects. Only a small fraction of X-linked genes showed sex-biased expression without functional consequences (4.0%, including *PRKX*) or no detectable sex effects (3.0%). No genes showed unclear mediation despite a sex difference or opposite-sign total and direct effects.

Together, these analyses demonstrate that while sex-biased gene expression can contribute to sex-biased gene essentiality, most sex differences in essentiality arise through expression-independent mechanisms, particularly for X-linked genes.

## Discussion

In this study, we investigated how sex chromosome composition relates to gene expression and gene essentiality across human cancer cell lines. Across the genome, we find that sex-biased gene expression can contribute to sex-biased gene essentiality, but this relationship is generally modest and not the dominant driver of sex differences in functional dependency. Instead, sex-biased essentiality is most often associated with expression-independent effects, particularly for genes on the X chromosome.

Variance partitioning showed that gene expression and sex chromosome dosage each explain only a small fraction of the overall variance in gene essentiality, with lineage accounting for a larger component and most variance remaining unexplained. These results indicate that while expression and dosage have measurable effects, gene essentiality is shaped by a broader set of factors not captured by baseline transcript abundance or chromosome copy number alone.

Consistent with this, genome-wide comparisons revealed that sex effects on gene expression and essentiality are small in magnitude but weakly aligned in direction. Genes that are more highly expressed in one sex tend, on average, to also be more essential in that sex, but this relationship is subtle and varies across genomic contexts. In particular, this alignment is limited among autosomal genes and more apparent for testis-elevated genes. These findings suggest that while expression differences may bias functional dependency in a consistent direction, they are not sufficient to explain most sex differences in essentiality.

Our mediation analyses further clarify the relationship between expression and essentiality by decomposing sex effects into expression-mediated and expression-independent components. Across genes, most sex differences in essentiality were associated with direct effects that are not explained by expression, whereas relatively few genes showed evidence consistent with expression-mediated effects. These results suggest that baseline transcriptional differences alone rarely account for sex-biased functional dependency.

Importantly, the balance between expression-mediated and expression-independent effects differs across genomic contexts. On autosomes, the most common pattern was a direct sex effect on essentiality without evidence of mediation, alongside a similarly large set of genes showing sex-biased expression without corresponding functional consequences. This indicates that transcriptional sex differences frequently do not translate into differences in cellular dependency. Expression-mediated effects were present for a subset of genes, and cases of inconsistent mediation – where direct and mediated effects act in opposite directions – suggest the presence of compensatory or buffering mechanisms.

In contrast, X-linked genes were strongly enriched for direct, expression-independent effects of sex on gene essentiality. This pattern is consistent with categorical sex differences in the X chromosome dosage differences and with the unique biology of the X chromosome, incomplete X inactivation and functional compensation between X-Y gametologs[28,29]. At the same time, expression-mediated effects were somewhat more frequent on the X chromosome than on autosomes, particularly among the X gametologs, suggesting that dosage-sensitive expression can contribute to sex-biased dependency in specific cases. Together, these results indicate that sex chromosome biology shapes functional dependency through mechanisms that are only partially captured by gene expression levels.

More broadly, our findings refine the role of sex-biased gene expression in shaping sex differences in cellular phenotypes. While prior work has emphasized transcriptional differences as a potential driver of sex-biased disease risk [4–6], our results suggest that such differences do not generally translate directly into differences in gene essentiality. Instead, sex-biased functional dependency appears to arise from a combination of mechanisms, with expression-mediated effects representing one component within a larger framework of sex-specific cellular regulation.

Several limitations should be considered. First, our analyses are based on cancer cell lines, which may not fully recapitulate the regulatory environments of primary tissues. Second, mediation analyses in this context identify statistical patterns consistent with indirect effects but do not establish causal mechanisms, and they do not account for additional layers of regulation such as post-transcriptional processes or protein-level effects. Third, gene sets used for targeted analyses may be incomplete or heterogeneous, particularly for complex diseases.

Despite these limitations, our study provides a systematic framework for linking sex-biased expression to functional dependency at a genome-wide scale. By integrating gene expression and CRISPR-based essentiality data, we show that sex-biased expression can contribute to sex differences in gene essentiality, but that most sex-biased functional effects are likely driven by expression-independent mechanisms, particularly in the context of sex chromosome biology.

## Acknowledgements

This work was supported by the Intramural Research Program of the National Institute on Aging. The contributions of the NIH authors are considered Works of the United States Government. The findings and conclusions presented in this paper are those of the authors and do not necessarily reflect the views of the NIH or the U.S. Department of Health and Human Services.

## Supplementary Table legends

**Supplementary Table 1. Summary of correlation and concordance analyses between Expression FC and Essentiality FC across gene sets and contrasts**. Each row represents a specific analysis defined by dataset, contrast, fold-change type (signed or absolute), gene set (all, autosomal, X chromosome, XCI escapees), and FDR threshold. For each combination, the number of genes included (n genes), Pearson’s correlation coefficient (r) and p-value, Spearman’s rank correlation coefficient (ρ) and p-value, the percentage of genes showing concordant fold-change direction, and results from a binomial test assessing concordance significance are reported. “Either < 0.2” indicates that genes were included if either the expression or essentiality FDR was below 0.2; “Expr and Ess < 0.2” indicates that threshold was met in both cases.

**Supplementary Table 2. Gene sets**. Each row represents a unique gene symbol, while each column corresponds to one gene list. Each cell indicates whether a given gene is present in the corresponding gene list, with values encoded as TRUE (gene present) or FALSE (gene absent).

**Supplementary Table 3. Variance partitioning of gene essentiality**. Gene-level variance partitioning results showing the proportion of variance in gene essentiality explained by gene expression, X chromosome dosage, Y chromosome dosage, sex chromosome complement, cell lineage, and residuals (unexplained variance). Each row corresponds to a gene, and values sum to 1 across columns.

**Supplementary Table 4. Gene-level mediation analysis of sex effects on gene essentiality**. Gene-level mediation results across three contrasts: XX vs. XY, X effect (XX vs. X0), and Y effect (Y+ vs. Y-). Columns report gene name, chromosome, contrast, and sample size (n_samples). Columns are grouped as follows: effects of sex chromosomes on gene expression (contrast → expression); effects of sex chromosomes on gene essentiality (contrast → essentiality); and effects of gene expression on essentiality (expression → essentiality), each reported as regression coefficients, standard errors (se), p-values, and FDR-adjusted p-values (p-adjusted). Mediation statistics include the average causal mediation effect (ACME), average direct effect (ADE), total effect, and proportion mediated, each reported with estimates, 95% confidence intervals, p-values, and FDR-adjusted p-values. Model fit statistics are reported as R^2^ values for the linear models used in the mediation framework: the expression model (expression ∼ contrast) and the essentiality model (essentiality ∼ contrast + expression).

**Supplementary Table 5. Enrichment of X-linked genes**. Results of Fisher’s exact tests assessing enrichment of X-linked genes across gene groups and expression-essentiality effect categories for three contrasts: XX vs. XY, X effect, and Y effect. Gene groups include total effect (TE), average direct effect (ADE), and average causal mediation effect (ACME) genes, as well as categories defined by the direction of sex-biased expression and essentiality. Columns report the odds ratio (OR), 95% confidence interval (CI), and p-value for enrichment.

**References and Notes**

Note 1: In this work, we use the term “sex” to refer to the classification assigned at birth based on anatomical characteristics, which often corresponds to common sex chromosome complements (XY individuals with testes and XX individuals with ovaries). We acknowledge that not all individuals fall within these typical categories, as variations exist in sex chromosomes, hormone levels and signaling, and physical traits. By “sex differences,” we refer to average differences observed between groups defined by these typical anatomical and chromosomal profiles. Furthermore, we distinguish sex from gender, which is a socially and culturally shaped and fluid concept. An individual’s gender identity may differ from their assigned sex, and because lived experiences are influenced by both biological and social factors, it is often difficult to fully separate these influences in human studies.

For our analyses, we used the sex annotation provided in the source datasets, corresponding to the reported sex of the tissue donor. These annotations are likely to reflect self-identified or clinically recorded sex and are not necessarily based on direct verification of sex chromosome complement. Consistent with this, available karyotype information indicates variability beyond typical XX and XY configurations (i.e., male samples with X0, XX, XXY, or XY; female samples with X0 or XX). Accordingly, our categorization reflects donor-reported sex rather than strictly chromosomal sex, and we interpret “sex differences” within this context. To account for inter-individual variation in sex chromosome complement, we additionally perform analyses based on X and Y chromosome dosage to assess potential differential effects.

